## Supporting Information 1 for "resevol: an R package for spatially explicit models of pesticide resistance given evolving pest genomes"

The evolutionary algorithm (Supporting Information 1)

A. Bradley Duthie[1 3], Rosie Mangan[1], Chintamani Rose McKeon[1], Matthew C. Tinsley[1], and Luc F. Bussière[2]

[1] Biological and Environmental Sciences, University of Stirling, Stirling, UK [2], Biological and Environmental Sciences 3A149 University of Stirling Stirling, FK9 4LA, UK

#### 1 Introduction

The resevol package models individuals with complex genomes that can include any number of haploid or diploid loci and any number of traits with an arbitrary pre-specified covariance structure. It does this by using a complex network mapping a path from the allele values at each loci to the covarying trait values of individuals.

The objective of the evolutionary algorithm in resevol is to find a set of network values that produce traits (red diamonds above) with the pre-specified covariance structure given allele values (green circles above) that are sampled from a standard normal distribution (i.e., mean of zero and standard deviation of one). Conceptually, the problem is simple; we need to find values for the black arrows above that produce traits that covary in the way that we want them to covary (within some acceptable margin of error). The `mine_gmatrix` function uses an evolutionary algorithm for identifying sets of values that work. An evolutionary algorithm is an algorithm that works especially well for *I know it when I see it* problems (Hamblin, 2013). Luke (2013) explains the idea behind these algorithms in more detail:

> They're algorithms used to find answers to problems when you have very little to help you: you don't know beforehand what the optimal solution looks like, you don't know how to go about finding it in a principled way, you have very little heuristic information to go on, and brute-force search is out of the question because the space is too large. But if you're given a candidate solution to your problem, you can test it and assess how good it is. That is, you know a good one when you see it.

In the `mine_gmatrix` function of the resevol package, an evolving population of networks like that in Figure 1 above is initialised. Parent networks produce offspring networks with some recombination and mutation, and offspring networks are selected based on how close their trait covariance matrix is to the pre-specified matrix input into the function. The algorithm is inspired by a similar algorithm within the GMSE R package (Duthie et al., 2018).

#### 2 Key data structures used

In the code, the arrows in the above Figure 1 are represented by a set of matrices that map loci values to trait values. There are 12 loci in Figure 1, and 4 nodes in each hidden layer (blue squares). Arrow values

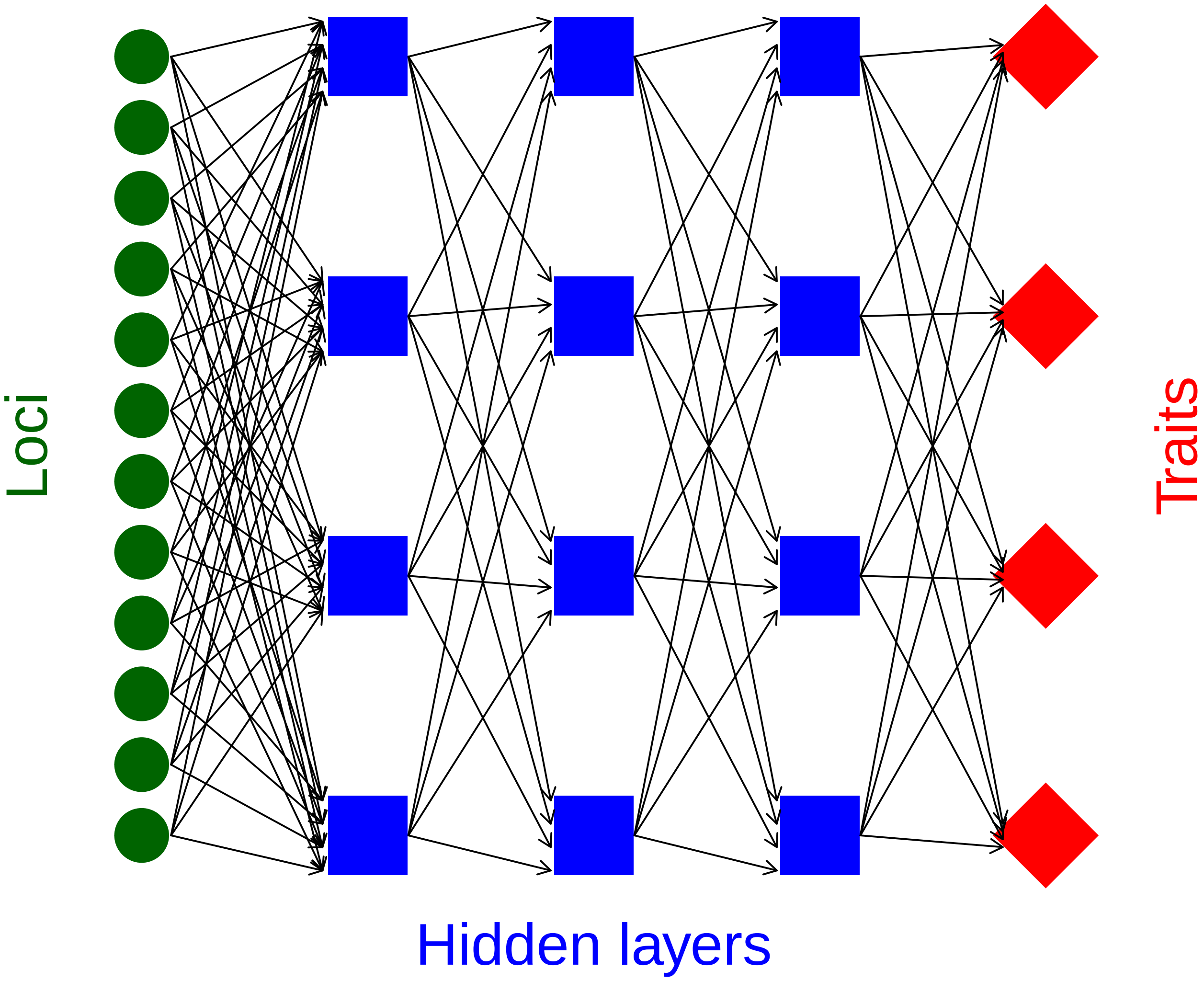


Figure 1: *Network mapping loci to traits through an intermediate set of hidden layers in the mine_gmatrix function*

between loci and the first hidden layer can then be represented by a matrix with 12 rows and 4 columns (i.e., row 1 holds a value for each of 4 arrows that point to the 4 hidden layer nodes). Note that the values below are initialised randomly, which is how they are initialised in the evolutionary algorithm.

```
arrows_1_dat <- rnorm(n = 4 * 12, mean = 0, sd = 0.1);
arrows_1_mat <- matrix(data = arrows_1_dat, nrow = 12, ncol = 4);
print(arrows_1_mat);
```

```
##               [,1]         [,2]        [,3]         [,4]
##  [1,] -0.047475891 -0.042684815  0.01290051 -0.011349773
##  [2,]  0.003719595 -0.088689293 -0.03180276  0.020113089
##  [3,] -0.105405274  0.095037211  0.02629605 -0.006940716
##  [4,] -0.104934618  0.136572718  0.13679253 -0.034114310
##  [5,] -0.042969241 -0.003485454  0.04502295 -0.068402422
##  [6,] -0.012586209 -0.053057403 -0.06232217 -0.078134223
##  [7,]  0.080142997  0.036078704  0.02414555 -0.071392556
##  [8,]  0.101615855  0.046973184  0.01945108  0.079247819
##  [9,] -0.137327773 -0.091940310 -0.04451465 -0.062169868
## [10,] -0.043598034  0.009897226  0.14991111 -0.005129893
## [11,] -0.040369246 -0.016800291 -0.12911360  0.007293362
## [12,]  0.007786802  0.093626656  0.14486569  0.024113093
```

We can initialise 12 allele values, one for each locus.

```
loci <- rnorm(n = 12, mean = 0, sd = 1);
```

To then get the value of the first column of four hidden layer nodes (i.e., the first column of blue squares in Figure 1), we can use matrix multiplication.

```
print(loci %*% arrows_1_mat);
```

```
##            [,1]       [,2]       [,3]       [,4]
## [1,] -0.3640695 0.01045695 -0.3796402 -0.2283317
```

We can likewise use a 4 by 4 square matrix to represent the values of the arrows from the first column of four hidden layer nodes to the second column of hidden layer nodes.

```
arrows_2_dat <- rnorm(n = 4 * 4, mean = 0, sd = 0.1);
arrows_2_mat <- matrix(data = arrows_2_dat, nrow = 4, ncol = 4);
print(arrows_2_mat);
```

```
##             [,1]         [,2]       [,3]         [,4]
## [1,]  0.11051532  0.001443321 0.05243207  0.007335664
## [2,] -0.07602310  0.052207800 0.18374055 -0.019406289
## [3,]  0.15501033  0.186791739 0.01750624 -0.193841413
## [4,] -0.01876937 -0.181548976 0.07359627 -0.001004679
```

We can then use matrix multiplication to map the 12 allele values to the values of the second column of hidden layer nodes.

```
print(loci %*% arrows_1_mat %*% arrows_2_mat);
```

```
##               [,1]        [,2]        [,3]       [,4]
## [1,] -0.09559273 -0.02943979 -0.04061799 0.07094577
```

This pattern can continue, with 4 by 4 square matrices representing the value of arrows between columns of hidden layer nodes, and between the last hidden layer column and traits (note that the number of hidden layer columns can be any natural number, but the number of nodes within a column always equals the number of traits). In the actual evolutionary algorithm code, all of these square matrices are themselves held in a large 3D array. But the idea is the same; a vector of allele values is multiplied by multiple matrices until a set of trait values is produced. If multiple vectors of random standard normal allele values are generated, then the traits that they produce from all of this matrix multiplication can be made to covary in some pre-specified way using the evolutionary algorithm.

### 3 General overview of the evolutionary algorithm

The evolutionary algorithm first initialises a population of networks, with each network having a unique set of values (i.e., black arrows in Figure 1, represented in the code by matrices explained in the previous selction). In each iteration of the evolutionary algorithm, with some probability, networks crossover their values with another randomly selected network. Individual values in each network then mutate with some probability. The fitness of each network is then calculated by comparing its trait covariances with those of a pre-specified covariance matrix. A tournament is then used to select the highest fitness networks, and those selected networks replace the old to comprise the new population. Iterations continue until either a maximum number of iterations is reached or a network is found that produces trait covariances sufficiently close to the pre-specified covariance matrix. Figure 2 below provides a general overview of the evolutionary algorithm.

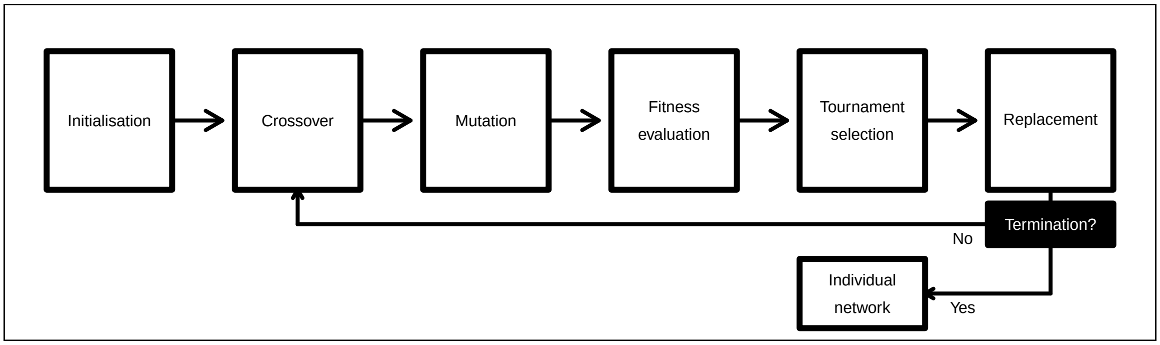


Figure 2: *Conceptual overview of the evolutionary algorithm used in the resevol package.*

The steps listed in the box above are explained in more detail below with reference to the arguments applied in the `mine_gmatrix` function that calls it.

#### 3.1 Initialisation

At the start of the evolutionary algorithm, a population of `npsize` networks is initialised. Each individual network in this population is represented by one matrix and one three dimensional array (see an explanation of the data structures above). All the elements of networks are initialised with a random sample from a normal distribution with a mean of 0 and a standard deviation of `sd_ini`.

#### 3.2 Crossover

An iteration of the evolutionary algorithm begins with a crossover between the values of networks in the population. For each network in the population, a crossover event will occur in the values linking loci to the first hidden layer (see Figure 1) with a probability of `pr_cross`. And a second independent crossover event will occur in the values linking the first hidden layer to traits, also with a probability of `pr_cross`. The reason that these two crossover events are independent is due to the different dimensions of the underlying arrays (see key data structures used above). A matrix with `loci` rows and a number of columns that matches the number of traits holds the values linking loci to the first hidden layer. A three dimensional array with row and column numbers matching trait number, and a depth matching the number of hidden `layers` (e.g., 3 in Figure 1) holds the remaining values linking the first hidden layer to trait values.

If a crossover event occurs for a focal network, then a contiguous set of values is defined and swapped with another randomly selected network in the population. Dimensions of the contiguous set are selected from a random uniform distribution. For example, given that the network in Figure 1 would be represented by a three dimensional array with 4 rows, 4 columns, and 3 layers, three random integers from 1-4, 1-4, and 1-3 would be sampled twice, respectively, with replacement. If the values selected were 1, 3, and 2 in the first sample, then 3, 3, and 1 in the second sample, then all values from rows 1-3, column 3, and layers 1-2 would be swapped between networks. Conceptually, this is the equivalent of drawing a square around set of arrows in Figure 1 and swapping the arrow values with the values of another network.

#### 3.3 Mutation

After crossover occurs, all network values mutate independently at a fixed probability of `mu_pr`. If a mutation event occurs, then a new value is randomly sampled from a normal distribution with a mean of 0 and a standard deviation of `mu_sd`. This value is then added to the existing value in the network.

#### 3.4 Fitness evaluation

After mutation, the fitness of each network in the population is evaluated. For each network, a set of `indivs` loci vectors is created, which represents the allele values of `indivs` potential individuals in a simulated population. Elements of each loci vector are randomly sampled from a standard normal distribution. For example, in the network of Figure 1 where `loci = 12`, `loci * indivs` standard normal values would be generated in total. After these `indivs` loci vectors are initialised, values in each vector are mapped to traits using the focal network, thereby producing `indivs` sets of traits from the focal network. These `indivs` sets of traits are then used to calculate the among trait covariances and build a trait covariance matrix for the network. This trait covariance matrix is then compared to the pre-specified `gmatrix` by calculating stress as the mean squared deviation between matrix elements. Lower stress values correspond to higher network fitness.

The stress of each network in the population of `npsize` networks is calculated using the above algorithm. Selection of the next generation of `npsize` networks is then done using a tournament.

#### 3.5 Tournament selection

After fitness evaluation, networks in the population compete in a series of tournaments to determine the composition of the next generation of `npsize` networks. Tournament selection is a flexible way to choose the fittest subset of the population (Hamblin, 2013). It starts by randomly selecting `sampleK` networks with replacement to form the competitors in a tournament (note that `sampleK` is constrained to be less than or equal to `npsize`). Of those `sampleK` networks, the `chooseK` networks with the highest fitness (i.e., lowest stress) are set aside to be placed in the new population (note that `chooseK` must be less than or equal to `sampleK`). More tournaments continue until a total of `npsize` new networks are set aside to form the new generation of networks.

##### 3.6 Termination

Throughout the evolutionary algorithm, the network with the lowest overall stress (from any generation) is retained. The evolutionary algorithm terminates if either the logged stress of the mean network is less than or equal to `term_cri`, or if `max_gen` generations of the evolutionary algorithm have occurred. The mean network stress is used instead of the lowest overall stress because error in randomly generated loci can result in unusually low stress values due to chance, which might not be replicated with a new random sample of loci. When the evolutionary algorithm terminates, only the network with the lowest overall stress is returned.

#### 4 Haploid and diploid individuals

The evolutionary algorithm does not distinguish between haploid and diploid genomes Instead, haploid and diploid individuals in resevol simulations are build differently from the mined network described above. For haploid individuals, network values are placed in individual genomes exactly as they are returned by `mine_gmatrix`. Hence, standard normal allele values at loci from haploid genomes will map to predictably covarying traits. For diploid individuals, all network values returned by `mine_gmatrix` are divided by 2, and two copies of each value are then placed in each individual to model diploid genomes. Allele values are then randomly sampled from a normal distribution with a mean of 0 and a standard deviation of `1 / sqrt(2)`, so that summed allele values at homologous loci will have a standard normal distribution. As such, effects of each loci are determined by the sum of homologous alleles. Similarly, homologous network values mapping allele values to traits are also summed, thereby producing the expected trait covariance structure.
