## Supporting Information 2 for "resevol: an R package for spatially explicit models of pesticide resistance given evolving pest genomes"

Advanced techniques (Supporting Information 2)

A. Bradley Duthie[1 3], Rosie Mangan[1], Chintamani Rose McKeon[1], Matthew C. Tinsley[1], and Luc F. Bussière[2]

[1] Biological and Environmental Sciences, University of Stirling, Stirling, UK [2], Biological and Environmental Sciences 3A149 University of Stirling Stirling, FK9 4LA, UK

#### 1 Introduction

Here we focus on advanced techniques for simulating pest ecological and evolutionary dynamics using the resevol R package. In the main text, we provided a simple example of individual-based simulations to demonstrate how to get started. This document focuses instead on demonstrating the more advanced options of the package and showcasing what it can do. The case study will focus on crop and pesticide rotation on a complex, customised landscape that includes farmland, grassland, forest, and water. The pest species will be sexually reproducing and obligately biparental, and its life history will include an egg and larval stage during which it feeds on crops and consumes pesticide, and a stage during which it moves, mates, and reproduces. The pest species will include a total of four evolving traits that affect the consumption of two crops and two pesticides. The objective of simulations will be to test how pest population size changes and traits evolve given different pesticide rotation regimes. Below, we explain how to model this system in detail, including all of the necessary code for reproducing the example within the text.

#### 2 Initialising pest genomes

First, we need to use the `mine_gmatrix` function to initialise pest genomes. In our example, individuals will have four covarying traits resulting from 12 loci. There will be 4 internal nodes that map loci values to traits (see the Evolutionary Algorithm explanation for details). The trait covariance structure will be defined as below.

```
library("resevol");
gmt <- matrix(data = c(1.0, -0.5, 0.2, 0.2, -0.5, 1.0, 0.2, 0.2, 0.2,
                       0.2, 0.4, -0.6, 0.2, 0.2, -0.6, 0.4), nrow = 4);
print(gmt);
```

```
##      [,1] [,2] [,3] [,4]
## [1,]  1.0 -0.5  0.2  0.2
## [2,] -0.5  1.0  0.2  0.2
## [3,]  0.2  0.2  0.4 -0.6
## [4,]  0.2  0.2 -0.6  0.4
```

Rows and columns 1 and 2 will represent traits underlying the consumption rate of crops 1 and 2, respectively. Hence, the variation in crop consumption rate is 1 for both crops, while the covariance in consumption rate is -0.5, meaning that there is a trade-off between pest ability to consume crop 1 versus crop 2. Rows and columns 3 and 4 will represent traits underlying the consumption rate of pesticides 1 and 2, respectively. Hence, the variation in pesticide consumption rate is 0.4 for both pesticides, which is lower than what it is for crops. There is a trade-off in the consumption rate of pesticide 1 versus pesticide 2, which is reflected in the covariance of -0.6 in the above matrix. Finally, there is a positive covariance of 0.2 between all crop and all pesticide consumption rates. What this means is that pests that consume crops quickly also tend to consume pesticides quickly, causing a potential trade-off between the beneficial effects of feeding ability and the negative effects of pesticides. We can now use the `mine_gmatrix` function to find a network mapping loci to traits that satisfies the above trait covariance structure. The options used below in `mine_gmatrix` arguments will lead to a computationally intense (and therefore time-consuming) search, particularly due to the large number of networks in the population (`npsize = 12000`), high number of individuals used to test network stress (`indivs = 2000`), high number of hidden nodes (`layers = 4`), strict stress criteria (`term_cri = -8`), and high maximum generation number (`max_gen = 5400`).

```
set.seed(2022);
mg  <- mine_gmatrix(gmatrix = gmt, loci = 12, indivs = 2000, npsize = 12000,
                    max_gen = 5400, sampleK = 1200, chooseK = 6, layers = 4,
                    mu_pr = 0.2, pr_cross = 0.2, mu_sd = 0.004,
                    term_cri = -8);
```

To save time, a genome `mg` produced from the code above has been saved into the resevol R package as `advanced_techniques_eg.rda`, which we can load. Note that running `mine_gmatrix` can be a time-consuming process, and will often require multiple attempts to get the above parameter settings to produce a desired network. See Hamblin (2013) for useful advice on parameter value selection for the evolutionary algorithm. Time-sensitive parameter values specific to the resevol package include `indivs` and `layers`. Increasing values for both of these parameters can increase run time.

```
load(system.file("advanced_eg.rda", package = "resevol"));
```

The contents of each list element of `mg` are not important for our purposes, but an explanation is available in the package documentation. What is important is the sixth list element, which holds the estimated covariance structure found by the evolutionary algorithm.

```
##            [,1]       [,2]       [,3]       [,4]
## [1,]  1.0677123 -0.4510309  0.1225596  0.1277905
## [2,] -0.4510309  1.0449414  0.1251180  0.1217368
## [3,]  0.1225596  0.1251180  0.5014936 -0.3992066
## [4,]  0.1277905  0.1217368 -0.3992066  0.5011398
```

We can compare the above covariance structure with the one that we specified in `gmt`. The stress of `mg` (i.e., mean squared deviation of `mg` elements from `gmt` elements) is 0.0099034. But this is only based on one population of `indivs = 2000` initialised pests. We can use the `stress_test` function to see what the distribution of stress is over 1000 such initialised populations of 2000 individuals (Figure 1).

```
sim_stress <- stress_test(mine_output = mg, indivs = 2000, reps = 1000);
hist(x = sim_stress, main = "", breaks = 20, xlab = "Initialised stress");
```

If we are satisfied with the distribution of stress, then we can accept `mg` as the genome for our initialised pests. The whole contents of `mg` will then be passed on to the `run_farm_sim` function, which initialises pests using the genome values in `mg` before running simulations. Before doing any of this, because we want to run a simulation on a customised landscape, we will explain how such a landscape can be generated and inserted into the simulation.

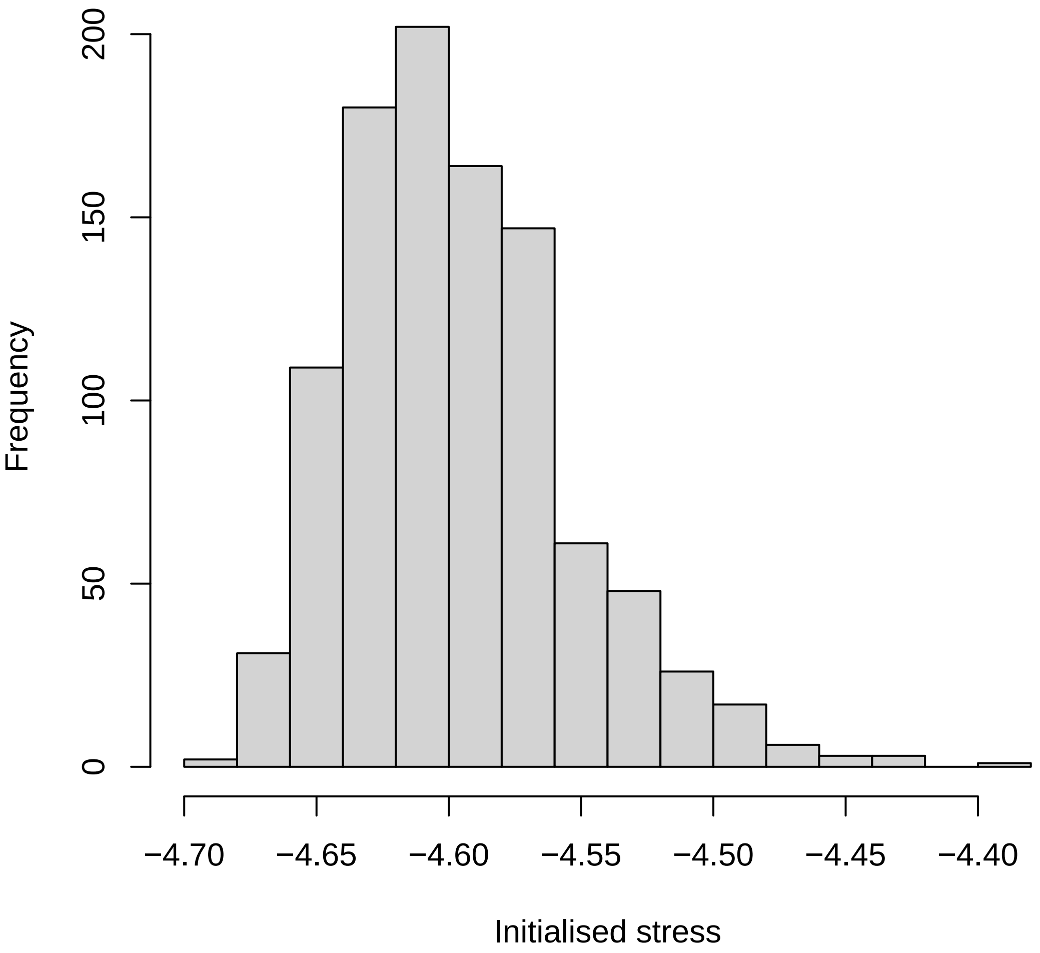


Figure 1: Distribution of stress for initialised pest trait covariances in the resevol R package. Stress values are computed using genome values produced by the mine_gmatrix function for 1000 replicate populations of initialised pest loci values.

### 3 Landscape

Default settings for running simulations with the `run_farm_sim` function create a spatially explicitly landscape with dimensions specified by arguments `xdim` and `ydim`, and a number of farms specified by `farms`. These arguments can be over-ridden with a matrix that defines landscape cell identities, which is set in the `terrain` argument. A very simple custom 12 by 12 landscape might look something like the below.

```
simple_terrain               <- matrix(data = 0, nrow = 12, ncol = 12)
simple_terrain[1:2, 1:3]     <- 1;
simple_terrain[3:7, 1:2]     <- 1;
simple_terrain[8:12, 1:2]    <- 2;
simple_terrain[1:2, 6:9]     <- 3;
simple_terrain[3:6, 7:9]     <- 3;
simple_terrain[1:6, 10:12]   <- 4;
simple_terrain[10:12, 7]     <- 5;
simple_terrain[7:12, 8:12]   <- 5;
simple_terrain[3:12, 3:4]    <- 6;
simple_terrain[7:9, 5]       <- 6;
simple_terrain[1:2, 4:5]     <- 7;
simple_terrain[3:6, 5:6]     <- 7;
simple_terrain[7:9, 6:7]     <- 7;
simple_terrain[10:12, 5:6]   <- 7;
print(simple_terrain)
```

```
##       [,1] [,2] [,3] [,4] [,5] [,6] [,7] [,8] [,9] [,10] [,11] [,12]
##  [1,]    1    1    1    7    7    3    3    3    3     4     4     4
##  [2,]    1    1    1    7    7    3    3    3    3     4     4     4
##  [3,]    1    1    6    6    7    7    3    3    3     4     4     4
##  [4,]    1    1    6    6    7    7    3    3    3     4     4     4
##  [5,]    1    1    6    6    7    7    3    3    3     4     4     4
##  [6,]    1    1    6    6    7    7    3    3    3     4     4     4
##  [7,]    1    1    6    6    6    7    7    5    5     5     5     5
##  [8,]    2    2    6    6    6    7    7    5    5     5     5     5
##  [9,]    2    2    6    6    6    7    7    5    5     5     5     5
## [10,]    2    2    6    6    7    7    5    5    5     5     5     5
## [11,]    2    2    6    6    7    7    5    5    5     5     5     5
## [12,]    2    2    6    6    7    7    5    5    5     5     5     5
```

What is important is that the matrix elements only include natural numbers from 1 to the total number of farms (not skipping any numbers). In the case of the above, numbers 1-7 all appear on the landscape, so the matrix will work. We can visualise the matrix more clearly by looking at it as an image (Figure 2).

```
par(mar = c(0, 0, 0, 0));
image(t(simple_terrain), xaxt = "n", yaxt = "n", useRaster = TRUE);
```

In Figure 2, each colour represents a unique farm, and colours correspond to the numbers within the matrix `simple_terrain` (note that the orientation differs between the matrix and the image). This is a simple example, but matrices can be made large to model highly complex landscapes. Additionally, not every colour needs to represent a 'farm' per se. We can represent other types of terrain such as grassland, woodland, or water with its own natural number. The trick in this case is to have each of these other types of terrain grow a crop and use a pesticide that has no effect on pests (i.e., cannot be consumed). If terrain types have no effect on pests, then they can effectively model land that is not used for farming. We can show this using the

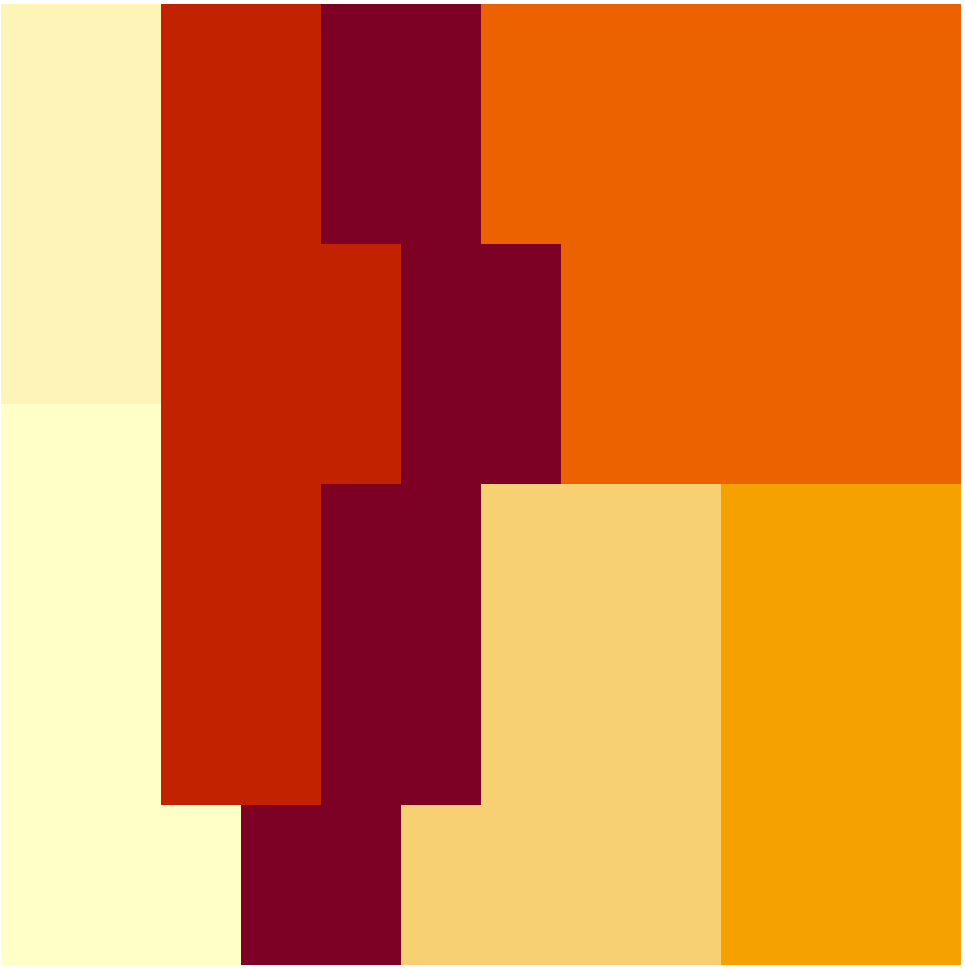

Figure 2: Image representing a simple landscape for the resevol package created from a 12 by 12 matrix of natural numbers from 1-7.

matrix `land_eg`, which is also included in the resevol package. The dimensions of the landscape modelled in `land_eg` are 128 by 128 cells, and the landscape includes 14 farms (1-14), grassland (15), forest (16), and water (17). The code below produces an image that shows how these terrain types are distributed over the landscape (Figure 3).

```
land_file <- system.file("landscape_eg.csv", package = "resevol");
land_dat  <- read.csv(file = land_file, header = FALSE);
land_eg   <- t(as.matrix(land_dat));
farm_cols <- c("#f4eadc", "#6a4b20", "#cea05f", "#e1c59d", "#a97833", "#cea05f",
               "#f2e6d6", "#6a4b20", "#cc9c59", "#dfc197", "#a27331", "#f0e3d0",
               "#5d421c", "#ca9852");
land_cols <- c(farm_cols, "#00ab41", "#234F1E", "#2832C2");
par(mar = c(0, 0, 0, 0));
image(land_eg, xaxt = "n", yaxt = "n", col = land_cols);
points(x = 0.2, y = 0.05, cex = 9, pch = 20);
text(x = 0.2, y = 0.05, labels = "1", cex = 2, col = "red");
points(x = 0.4, y = 0.1, cex = 9, pch = 20);
text(x = 0.4, y = 0.1, labels = "2", cex = 2, col = "red");
points(x = 0.4, y = 0.1, cex = 9, pch = 20);
text(x = 0.4, y = 0.1, labels = "2", cex = 2, col = "red");
points(x = 0.25, y = 0.27, cex = 9, pch = 20);
text(x = 0.25, y = 0.27, labels = "3", cex = 2, col = "red");
points(x = 0.20, y = 0.43, cex = 9, pch = 20);
text(x = 0.20, y = 0.43, labels = "4", cex = 2, col = "red");
points(x = 0.42, y = 0.48, cex = 9, pch = 20);
text(x = 0.42, y = 0.48, labels = "5", cex = 2, col = "red");
points(x = 0.28, y = 0.58, cex = 9, pch = 20);
text(x = 0.28, y = 0.58, labels = "6", cex = 2, col = "red");
points(x = 0.1, y = 0.8, cex = 9, pch = 20);
text(x = 0.1, y = 0.8, labels = "7", cex = 2, col = "red");
points(x = 0.7, y = 0.05, cex = 9, pch = 20);
text(x = 0.7, y = 0.05, labels = "8", cex = 2, col = "red");
points(x = 0.9, y = 0.2, cex = 9, pch = 20);
text(x = 0.9, y = 0.2, labels = "9", cex = 2, col = "red");
points(x = 0.85, y = 0.4, cex = 9, pch = 20);
text(x = 0.85, y = 0.4, labels = "10", cex = 2, col = "red");
points(x = 0.92, y = 0.58, cex = 9, pch = 20);
text(x = 0.92, y = 0.58, labels = "11", cex = 2, col = "red");
points(x = 0.88, y = 0.76, cex = 9, pch = 20);
text(x = 0.88, y = 0.76, labels = "12", cex = 2, col = "red");
points(x = 0.86, y = 0.93, cex = 9, pch = 20);
text(x = 0.86, y = 0.93, labels = "13", cex = 2, col = "red");
points(x = 0.52, y = 0.91, cex = 9, pch = 20);
text(x = 0.52, y = 0.91, labels = "14", cex = 2, col = "red");
points(x = 0.05, y = 0.59, cex = 7, pch = 20);
text(x = 0.05, y = 0.59, labels = "15", cex = 1.7, col = "white");
points(x = 0.46, y = 0.28, cex = 7, pch = 20);
text(x = 0.46, y = 0.28, labels = "16", cex = 1.7, col = "white");
points(x = 0.62, y = 0.36, cex = 7, pch = 20);
text(x = 0.62, y = 0.36, labels = "17", cex = 1.7, col = "white");
```

This `land_eg` matrix can be used in `run_farm_sim` by setting `terrain = land_eg`. The dimensions of the landscape will then automatically be set to `xdim = 128` and `ydim = 128`, and the number of farms will be set to `farms = 17`. To ensure that we do not have pests consuming crops or pesticides from non-farm

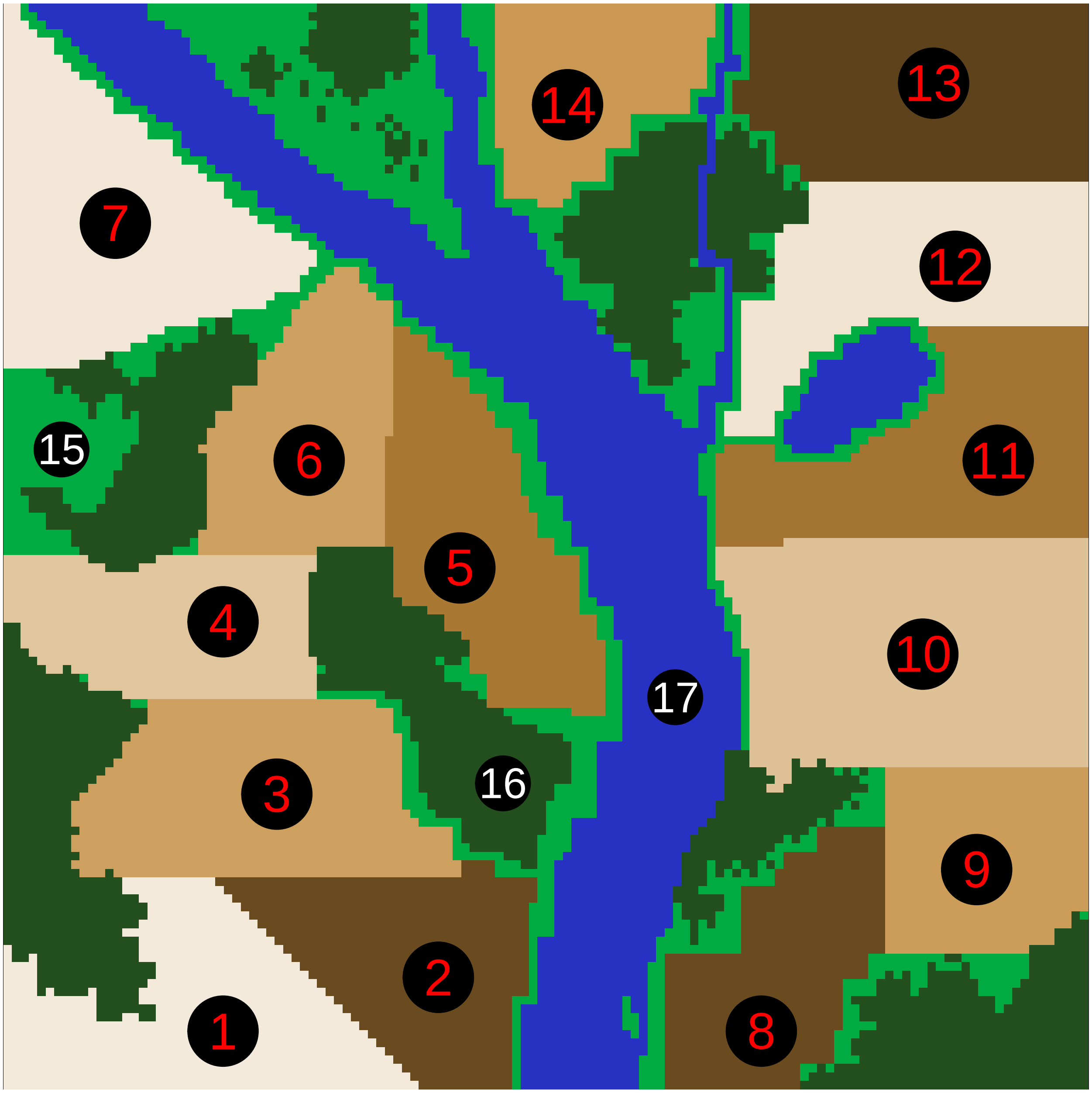


Figure 3: A complex landscape to be used in the resevol R package, including 14 separate farms, grassland, forest, and water. The image is represented in the code by a matrix in which element numbers correspond to different terrain colours (e.g., elements corresponding to water are numbered 17).

areas, we just need to set pest consumption options accordingly. Next, we describe how this can be done by defining how crops and pesticides will be initialised and rotated on the landscape.

### 4 Running simulations

We will simulate the same population twice using the `mg` pest genome and `land_eg` landscape described above under two different pesticide regimes, simulation 1 and simulation 2. In both regimes, the same pesticide is applied to farms that occupy the same side of the river (e.g., farms 1-7 apply pesticide 1 and farms 8-14 apply pesticide 2). In simulation 1, the pesticide is never rotated. In simulation 2, farms rotate between pesticides 1 and 2 every 9 time steps. First, we will demonstrate how to customise crop and pesticide rotation for simulation 1, then run the simulation, and finally show the output. Second, we will repeat the process for simulation 2 and demonstrate how comparisons can be made between the two simulations.

#### 4.1 Simulation 1

First, we focus on how to set custom pesticide and crop rotation for simulations. Next, we run simulation 1 with the `run_farm_sim` function. Finally, we demonstrate how to interpret output for simulation 1.

##### 4.1.1 Custom pesticide and crop rotation

The `run_farm_sim` arguments `crop_init` and `crop_rotation_type` specify starting crops for each farm and how crops rotate, respectively. Equivalent arguments `pesticide_init` and `pesticide_rotation_type` similarly specify starting pesticides and pesticide rotations, respectively. We can set each argument to a number as a short-hand way of specifying these parameters. For example, the default initialisation `crop_init = "random"` simply causes each farm to start the simulation with a random crop. Setting values of `crop_rotation_type` to 1, 2, or 3 causes no rotation, random rotation, or cycling through crop numbers in sequence, respectively. But we can generate more specific initialisation and rotation types using a vector to initialise crops and pesticides, and using a matrix to define rotation. For initialisation, vector elements correspond to farm number and element values correspond to the crop initialised. For our example with 17 unique zones of terrain on the landscape, we can define initialised crop choice as follows.

```
initial_crop <- c(1, 1, 1, 1, 1, 1, 1, 2, 2, 2, 2, 2, 2, 2, 3, 3, 3);
```

That is, the first 7 farms on one side of the river all start off growing crop 1. Farms 8-14 all start off growing crop 2. Crop 3 is used as a dummy for grassland (15), forest (16), and water (17). We explain how to ensure that 15-17 is never used by pests below. Meanwhile, we can specify a matrix for the crop rotation regime. This regime will cause farmers to rotate between crops 1 and 2, but zones 15-17 to always use 'crop' 3.

```
rotate_crop        <- matrix(data = 0, nrow = 3, ncol = 3);
rotate_crop[1, 2] <- 1;
rotate_crop[2, 1] <- 1;
rotate_crop[3, 3] <- 1;
print(rotate_crop);
```

```
##      [,1] [,2] [,3]
## [1,]    0    1    0
## [2,]    1    0    0
## [3,]    0    0    1
```

In the above `rotate_crop`, matrix elements define the probability of transitioning from one crop type (rows) to another crop type (columns). Hence, in `rotate_crop`, farms that have been applying crop 1 will always switch to crop 2, and vice versa. Landscape zones applying crop 3 will never transition to any other crop, nor will any other crop transition to type 3. We can set initial pesticides and pesticide rotation in the same way. First we set initial pesticide use.

```
initial_pesticide <- c(1, 1, 1, 1, 1, 1, 1, 2, 2, 2, 2, 2, 2, 2, 3, 3, 3);
```

Next, we define pesticide rotation. Recall that in simulation 1, pesticides will not rotate and farms will maintain the same pesticide type over all time steps.

```
rotate_pesticide           <- matrix(data = 0, nrow = 3, ncol = 3);
rotate_pesticide[1, 1]     <- 1;
rotate_pesticide[2, 2]     <- 1;
rotate_pesticide[3, 3]     <- 1;
print(rotate_pesticide);
```

```
##      [,1] [,2] [,3]
## [1,]    1    0    0
## [2,]    0    1    0
## [3,]    0    0    1
```

We therefore have set farms to always use one of two crops and one of two pesticides. Everything that is not a farm will use crop 3 and pesticide 3. Next, we bring everything together and show how pests, the landscape, and the rotation of crops and pesticides can be used in `run_farm_sim` for simulations.

##### 4.1.2 Running simulation 1

The code below runs simulation 1 using the `run_farm_sim` function.

```
set.seed(2022);
sim1       <- run_farm_sim(mine_output             = mg,
                           terrain                 = land_eg,
                           crop_init               = initial_crop,
                           crop_rotation_type      = rotate_crop,
                           pesticide_init          = initial_pesticide,
                           pesticide_rotation_type = rotate_pesticide,
                           food_consume            = c("T1", "T2", 0),
                           pesticide_consume       = c("T3", "T4", 0),
                           crop_number             = 3,
                           pesticide_number        = 3,
                           trait_means             = c(2, 2, 0.0, 0.0),
                           max_age                 = 6,
                           min_age_feed            = 0,
                           max_age_feed            = 2,
                           min_age_move            = 3,
                           max_age_move            = 6,
                           min_age_metabolism      = 3,
                           max_age_metabolism      = 6,
                           metabolism              = 0.5,
                           food_needed_surv        = 1,
                           reproduction_type       = "food_based",
```

```
                          food_needed_repr          = 2,
                          N                         = 1000,
                          repro                     = "biparental",
                          mating_distance           = 4,
                          rand_age                  = TRUE,
                          pesticide_tolerated_surv = 2,
                          movement_bouts            = 4,
                          move_distance             = 2,
                          crop_per_cell             = 10,
                          crop_sd                   = 0,
                          pesticide_per_cell        = 1,
                          pesticide_sd              = 0,
                          crop_rotation_time        = 18,
                          pesticide_rotation_time  = 9,
                          time_steps                = 240,
                          pesticide_start           = 81,
                          immigration_rate          = 100,
                          land_edge                 = "reflect",
                          mutation_pr               = 0.01,
                          crossover_pr              = 0.01,
                          print_gens               = TRUE,
                          print_last               = TRUE);
```

Immediately below the first six arguments, we have the arguments `food_consume`, `pesticide_consume`, `crop_number`, and `pesticide_number`. The way that these arguments are set ensures that pests only interact with farm cells. The rate of food consumption for the first two crops is defined by the values of traits 1 and 2 and set as `"T1"` and `"T2"`, respectively. The rate of consumption for crop 3 is set to 0, meaning that no food can be eaten on these cells. Similarly, the rate at which pesticides 1 and 2 are consumed is defined by the values of traits 3 and 4, which are set as `"T3"` and `"T4"`, respectively. The rate of consumption for pesticide 3 is set to 0 so that pesticide is not consumed on these cells.

We next set the mean values of traits at the start of the simulation with the argument `trait_means`. Mean consumption rate of both crops is set to 2, while mean pesticide consumption rate is set to 0 (note, negative consumption rate values are possible, but are realised as no consumption). Pests live up to 6 time steps (`max_age = 6`), and are defined to start feeding upon birth (`min_age_feed = 0`) and stop at age 2 (`max_age_feed = 2`). At age 3, pests start to move (`min_age_move = 3`) and metabolise food (`min_age_metabolism = 3`) at a rate of `metabolism = 0.5`. The `metabolism` argument defines the amount of consumed food lost during the time step, which can affect pests when food consumed affects survival (`food_needed_surv`) and reproduction (`reproduction_type`). By setting `reproduction_type = "food_based"` and `food_needed_surv = 1`, we model a system in which pests burn the food consumed from ages 0-2 at a rate of 0.5 units per time step after age 3, then die when their reserves drop below 1. By setting `food_needed_repr = 2`, we ensure that only pests that have 2 or more units of food stored can reproduce. Note that if we desired, we could also set `food_needed_surv`, `food_needed_repr`, and `metabolism` to be evolving traits.

We initialise the population with `N = 1000` obligately biparental (`repro = "biparental"`) pests, and can find a mate within `mating_distance = 4` cells of their own cell. All pests are initialised at a random age (`rand_age = TRUE`). Pests will die if they consume more than 2 total units of pesticide (`pesticide_tolerated_surv = 2`). In each time step, pests that are able to move will do so an average of 4 times on the landscape (`movement_bouts = 4`), moving up to 2 cells in any direction during each bout (`movement_distance = 2`). Recall that since pests consume food and pesticide only between ages 0-2, then start moving at age 3, no food or pesticide consumption occurs during movement (if applicable, it could be turned on with `feed_while_moving = TRUE` and `pesticide_while_moving = TRUE`). Each farm cell produces 10 units of crops (`crop_per_cell = 10` and `crop_sd = 0`) and 1 unit of pesticide (`pesticide_per_cell = 1` and `pesticide_sd = 0`). Crops are rotated every 18 time steps (`crop_rotation_time = 18`), and pesticides are set to rotate every 9 time steps (`pesticide_rotation_time = 9`), but this pesticide rotation has no effect in simulation 1 because the same pesticide is applied upon rotation (as defined by `rotate_pesticide`). This can be conceptualised as modelling an 18 time step growing season in which pesticides are re-applied at the start and halfway point of a season. Note that it is critical to consider that crops are therefore refreshed every 18 time steps, not every time step. Hence, any food consumed by a pest will be lost from a cell and not replenished until crop rotation occurs. Because more than one pest can occupy a single cell, crop loss can occur quickly depending on how parameter values are set.

Lastly, we set the number of time steps to `time_steps = 240`, and we set `pesticide_start = 81`, which means that pesticides are not applied at all until after a burn-in of 81 time steps. In each time step, an average of 100 immigrants enter the population (`immigration_rate = 100`). We set the landscape edge type to `land_edge = "reflect"` to model a reflective edge in which pests that attempt to leave one side of the landscape bounce back toward the centre. Pest genome mutation rate and crossover rate at a locus are set to `mutation_pr = 0.01` and `crossover_pr = 0.01`, respectively. We print out the dynamics of the evolving population over time, and all of the individual data from the last time step, to two separate CSV files (individual data are not included in the package due to its size). All arguments of `run_farm_sim` not mentioned are set to default parameter values, which can be found in the package documentation.

##### 4.1.3 Simulation 1 output

When running the function `run_farm_sim`, the population size of the pest will be printed in the R console in each time step. This is primarily because simulations can take a long time, and printing makes it possible to estimate how much time is remaining. Printing to the console can be turned off by setting `print_gens = FALSE`. Once the simulation has finished, two CSV files will be created in the working directory. The file "population_data.csv" saves population data over time, and "individuals.csv" saves every characteristic of all individuals in the last time step (including full individual genomes). Here we show how to work with the most relevant information from these two files to make inferences about pest population and evolutionary dynamics.

We will start with the population level output that is printed to "population_data.csv", which has been renamed "population_data_sim1.csv".

```
population_data_file_sim1 <- system.file("population_data_sim1.csv",
                                         package = "resevol");
population_data_sim1      <- read.csv(file = population_data_file_sim1);
```

This file includes population-level parameters of population size and mean pest age, sex, food consumed, pesticide consumed, mortality rate, and trait values reported for each time step. If `get_f_coef = TRUE` in `run_farm_sim`, then mean inbreeding coefficients of pests are also reported.

```
print(head(population_data_sim1));
```

```
##   time_step population_size mean_age mean_sex mean_food_consumed
## 1         0            1000 3.982000 2.499000           0.396658
## 2         1             301 1.813953 2.498339           2.590001
## 3         2             415 2.166265 2.474699           2.708252
## 4         3             540 2.412963 2.488889           2.830218
## 5         4             756 2.488095 2.469577           2.755487
## 6         5             964 2.452282 2.484440           2.819527
##   mean_pesticide_consumed mortality_rate mean_food1_consumed
## 1                       0       0.837000            0.179506
```

```
## 2                       0       0.205980           0.578690
## 3                       0       0.233735           0.417990
## 4                       0       0.218519           0.398647
## 5                       0       0.279101           0.411980
## 6                       0       0.312241           0.405805
##   mean_food2_consumed mean_food3_consumed mean_pesticide1_consumed
## 1            0.217152                   0                        0
## 2            0.738026                   0                        0
## 3            0.526348                   0                        0
## 4            0.521056                   0                        0
## 5            0.496357                   0                        0
## 6            0.468015                   0                        0
##   mean_pesticide2_consumed mean_pesticide3_consumed trait1_mean_value
## 1                        0                        0          1.998091
## 2                        0                        0          1.962117
## 3                        0                        0          2.021925
## 4                        0                        0          2.020423
## 5                        0                        0          2.073448
## 6                        0                        0          2.083852
##   trait2_mean_value trait3_mean_value trait4_mean_value mean_f
## 1          2.014113          0.001834          0.003205      0
## 2          2.140079          0.041814          0.000534      0
## 3          2.211915          0.067218          0.029188      0
## 4          2.279216          0.034372          0.089154      0
## 5          2.313421          0.099480          0.059904      0
## 6          2.367128          0.105447          0.080354      0
```

Because the structure of the data frame above depends on the number of crops, pesticides, and evolving traits simulated, there are no pre-set functions in the resevol package for plotting. Here we include code for plotting population size, consumption of the two food types, two pesticides types, and evolving trait means. Note that because crop and pesticide 3 are dummy variables (not consumed at all), we do not need to plot these. Figure 4 shows how population density of pests changes over time.

```
mbox <- function(x0, x1, y0, y1){
    xx <- seq(from=x0, to=x1, length.out = 100);
    yy <- seq(from=y0, to=y1, length.out = 100);
    xd <- c(rep(x0, 100), xx, rep(x1,100), rev(xx));
    yd <- c(yy, rep(y1,100), rev(yy), rep(y0, 100));
    return(list(x=xd, y=yd));
}
season          <- seq(from = 0, to = 240, by = 18);
blocks          <- length(season) - 1;
plot(x = population_data_sim1[["time_step"]], type = "n",
     y = population_data_sim1[["population_size"]], cex.lab = 1.25,
     cex.axis = 1.25, ylab = "Pest population abundance", xlab = "Time step",
     ylim = c(0, max(population_data_sim1[["population_size"]])));
for(i in 1:blocks){
    rbox <- mbox(x0 = season[i], x1 = season[i + 1], y0 = -1000, y1 = 21000);
    if(i %% 2 == 0){
        polygon(x = rbox$x, y = rbox$y, lwd = 3, border = NA,
                col = "grey90");
    }
}
points(x = population_data_sim1[["time_step"]],
```

```
     y = population_data_sim1[["population_size"]], type = "l", lwd = 2);
abline(v = 81, lwd = 3, col = "red");
box();
```

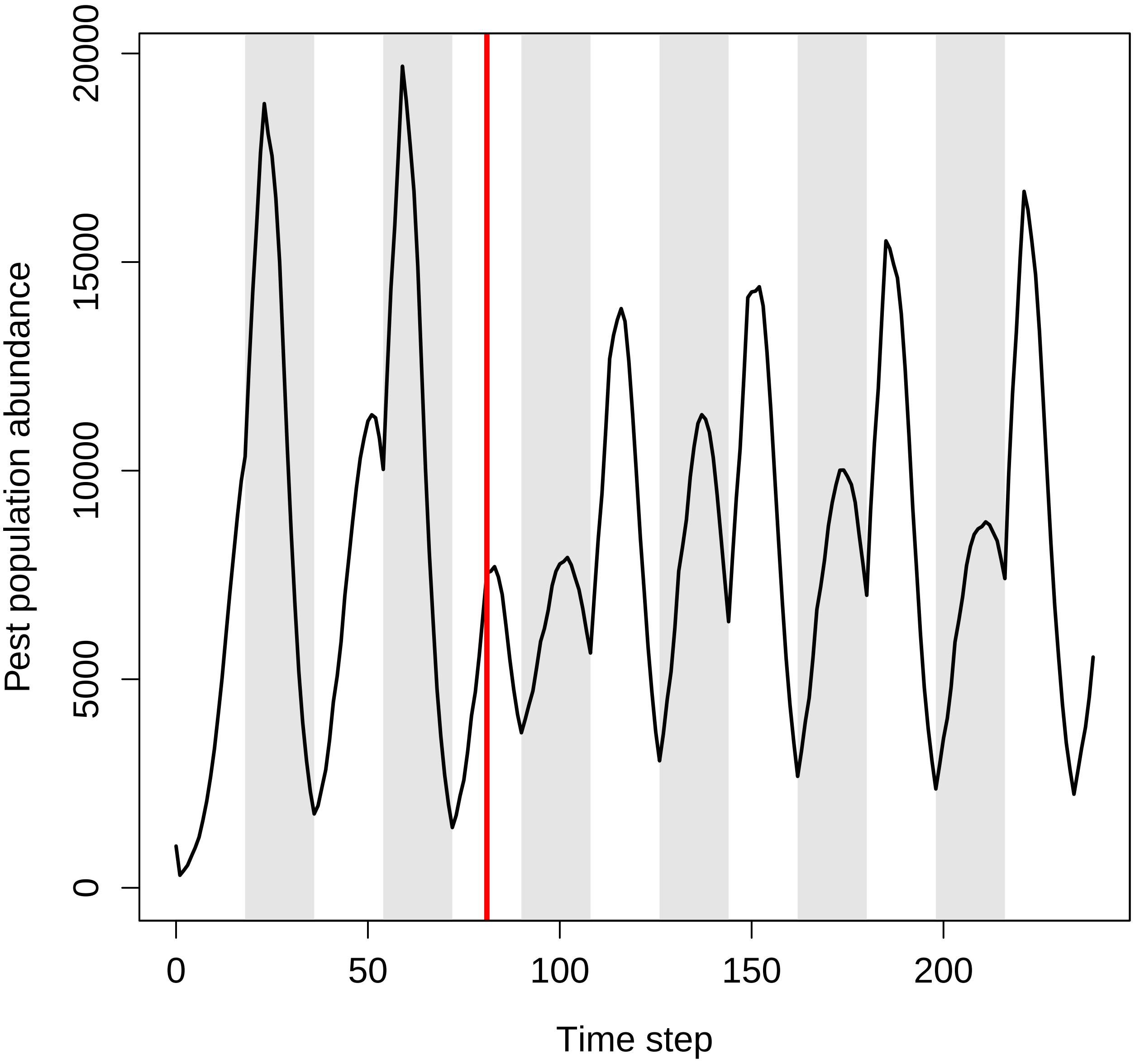


Figure 4: Pest abundance over time in a population simulated using the resevol R package in which pesticides are not rotated over time. The grey shaded regions show individual crop seasons; pesticides are applied at the start and midpoint of each seasons, beginning at the time step indicated by the red vertical line.

Figure 5 shows how the mean consumption of different foods and pesticides changes over time.

```
par(mfrow = c(2, 2), mar = c(1, 4, 1, 1));
plot(x = population_data_sim1[["time_step"]], type = "n", ylim = c(0, 0.8),
     y = population_data_sim1[["mean_food1_consumed"]], cex.lab = 1.25,
     ylab = "Mean crop 1 consumed", xlab = "Time step", xaxt = "n");
```

```
for(i in 1:blocks){
    rbox <- mbox(x0 = season[i], x1 = season[i + 1], y0 = -1000, y1 = 6000);
    if(i %% 2 == 0){
        polygon(x = rbox$x, y = rbox$y, lwd = 3, border = NA,
                col = "grey90");
    }
}
points(x = population_data_sim1[["time_step"]],
       y = population_data_sim1[["mean_food1_consumed"]], type = "l", lwd = 2);
text(x = 5, y = 0.79, labels = "a", cex = 2);
box();
par(mar = c(1, 4, 1, 1));
plot(x = population_data_sim1[["time_step"]], type = "n", ylim = c(0, 0.8),
     y = population_data_sim1[["mean_food2_consumed"]], cex.lab = 1.25 ,
     ylab = "Mean crop 2 consumed", xlab = "Time step", xaxt = "n");
for(i in 1:blocks){
    rbox <- mbox(x0 = season[i], x1 = season[i + 1], y0 = -1000, y1 = 6000);
    if(i %% 2 == 0){
        polygon(x = rbox$x, y = rbox$y, lwd = 3, border = NA,
                col = "grey90");
    }
}
points(x = population_data_sim1[["time_step"]],
       y = population_data_sim1[["mean_food2_consumed"]], type = "l", lwd = 2);
text(x = 5, y = 0.79, labels = "b", cex = 2);
box();
par(mar = c(4, 4, 1, 1));
plot(x = population_data_sim1[["time_step"]], type = "n", ylim = c(0, 0.8),
     y = population_data_sim1[["mean_pesticide1_consumed"]], cex.lab = 1.25,
     ylab = "Mean pesticide 1 consumed", xlab = "Time step");
for(i in 1:blocks){
    rbox <- mbox(x0 = season[i], x1 = season[i + 1], y0 = -1000, y1 = 6000);
    if(i %% 2 == 0){
        polygon(x = rbox$x, y = rbox$y, lwd = 3, border = NA,
                col = "grey90");
    }
}
points(x = population_data_sim1[["time_step"]],
       y = population_data_sim1[["mean_pesticide1_consumed"]], type = "l", lwd = 2);
box();
text(x = 5, y = 0.78, labels = "c", cex = 2);
par(mar = c(4, 4, 1, 1));
plot(x = population_data_sim1[["time_step"]], type = "n", ylim = c(0, 0.8),
     y = population_data_sim1[["mean_pesticide2_consumed"]], cex.lab = 1.25,
     ylab = "Mean pesticide 2 consumed", xlab = "Time step");
for(i in 1:blocks){
    rbox <- mbox(x0 = season[i], x1 = season[i + 1], y0 = -1000, y1 = 6000);
    if(i %% 2 == 0){
        polygon(x = rbox$x, y = rbox$y, lwd = 3, border = NA,
                col = "grey90");
    }
}
points(x = population_data_sim1[["time_step"]],
```

```
      y = population_data_sim1[["mean_pesticide2_consumed"]], type = "l",
      lwd = 2);
box();
text(x = 5, y = 0.78, labels = "d", cex = 2);
```

Similarly, we can visualise how all four traits evolve over time (Figure 6).

```
par(mfrow = c(2, 2), mar = c(1, 4, 1, 1));
plot(x = population_data_sim1[["time_step"]], type = "n", ylim = c(2, 3.6),
     y = population_data_sim1[["trait1_mean_value"]], cex.lab = 1.25,
     ylab = "Mean crop 1 consumption trait", xlab = "Time step", xaxt = "n");
for(i in 1:blocks){
    rbox <- mbox(x0 = season[i], x1 = season[i + 1], y0 = -1000, y1 = 6000);
    if(i %% 2 == 0){
        polygon(x = rbox$x, y = rbox$y, lwd = 3, border = NA,
                col = "grey90");
    }
}
points(x = population_data_sim1[["time_step"]],
     y = population_data_sim1[["trait1_mean_value"]], type = "l", lwd = 2);
text(x = 5, y = 3.56, labels = "a", cex = 2);
box();
par(mar = c(1, 4, 1, 1));
plot(x = population_data_sim1[["time_step"]], type = "n", ylim = c(2, 3.6),
     y = population_data_sim1[["trait2_mean_value"]], cex.lab = 1.25 ,
     ylab = "Mean crop 2 consumption trait", xlab = "Time step", xaxt = "n");
for(i in 1:blocks){
    rbox <- mbox(x0 = season[i], x1 = season[i + 1], y0 = -1000, y1 = 6000);
    if(i %% 2 == 0){
        polygon(x = rbox$x, y = rbox$y, lwd = 3, border = NA,
                col = "grey90");
    }
}
points(x = population_data_sim1[["time_step"]],
       y = population_data_sim1[["trait2_mean_value"]], type = "l", lwd = 2);
text(x = 5, y = 3.56, labels = "b", cex = 2);
box();
par(mar = c(4, 4, 1, 1));
plot(x = population_data_sim1[["time_step"]], type = "n", ylim = c(-0.8, 1.4),
     y = population_data_sim1[["trait3_mean_value"]], cex.lab = 1.25,
     ylab = "Mean pesticide 1 consumption trait", xlab = "Time step");
for(i in 1:blocks){
    rbox <- mbox(x0 = season[i], x1 = season[i + 1], y0 = -1000, y1 = 6000);
    if(i %% 2 == 0){
        polygon(x = rbox$x, y = rbox$y, lwd = 3, border = NA,
                col = "grey90");
    }
}
points(x = population_data_sim1[["time_step"]],
       y = population_data_sim1[["trait3_mean_value"]], type = "l", lwd = 2);
box();
text(x = 5, y = 1.34, labels = "c", cex = 2);
par(mar = c(4, 4, 1, 1));
```

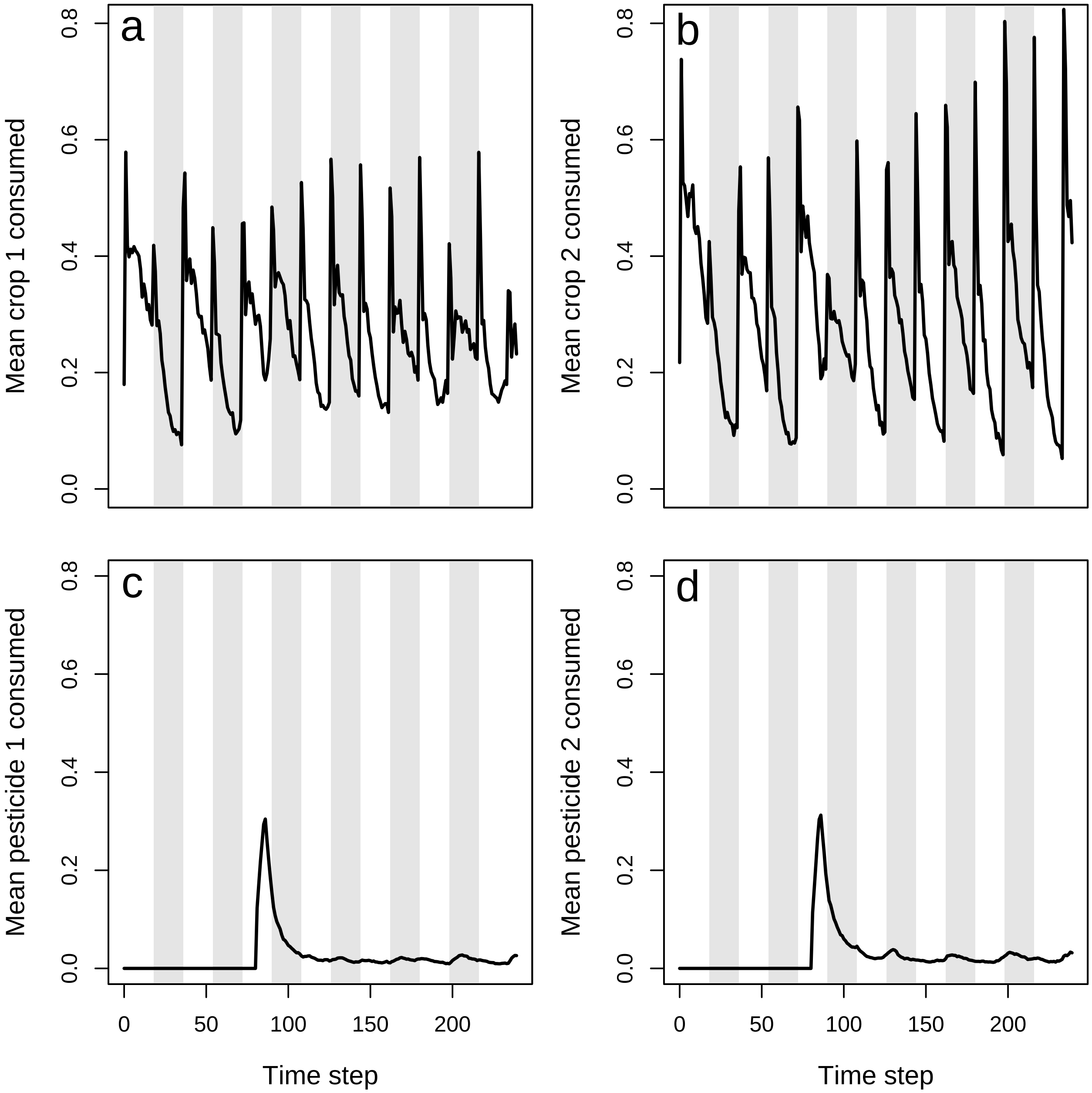


Figure 5: Pest consumption over time in a population simulated using the resevol R package in which pesticides are not rotated over time. Panels show (a) mean amount of crop 1 and (b) crop 2 consumed per pest, and (c) mean amount of pesticide 1 and (d) pesticide 2 consumed per pest. Grey shaded regions show individual crop seasons; pesticides are applied at the start and midpoint of each seasons, beginning at the time step 100.

```
plot(x = population_data_sim1[["time_step"]], type = "n", ylim = c(-0.8, 1.4),
     y = population_data_sim1[["trait4_mean_value"]], cex.lab = 1.25,
     ylab = "Mean pesticide 2 consumption trait", xlab = "Time step");
for(i in 1:blocks){
    rbox <- mbox(x0 = season[i], x1 = season[i + 1], y0 = -1000, y1 = 6000);
    if(i %% 2 == 0){
        polygon(x = rbox$x, y = rbox$y, lwd = 3, border = NA,
                col = "grey90");
    }
}
points(x = population_data_sim1[["time_step"]],
       y = population_data_sim1[["trait4_mean_value"]], type = "l", lwd = 2);
box();
text(x = 5, y = 1.34, labels = "d", cex = 2);
```

Next, we look at the file that includes the complete information for all individuals in the last time step of the simulation. Using this data set, we can examine the distribution of pests on the landscape, and the realised covariance of evolving pest traits. In this output "last_time_step.csv", traits always begin in column 102, meaning that individual trait values for traits 1, 2, 3, and 4 will be in columns 102, 103, 104, and 105, respectively. In this example, the CSV output file "last_time_step.csv" has been renamed "last_time_step_sim1.csv". To minimise the size of the resevol R package, all data columns except pest locations and traits have been removed from example outputs, so pest locations are in columns 1-2 (not 3-4) and traits are in columns 3-6 (not 102-105). The last time step output is read into R below.

```
last_time_step_file_sim1 <- system.file("last_time_step_sim1.csv",
                                        package = "resevol");
pop_last_time_step_sim1  <- read.csv(file = last_time_step_file_sim1);
```

We can plot the distribution over the landscape using the code below. The code identifies the x-locations and y-locations of each pest `i` in `pop_last_time_step_sim1`, and these landscape cells are replaced with the number 18. A new land colour for black `#000000` is then included, and the map is reproduced as an image in R. Figure 7 shows the same map as above, but with the numbering removed to make the pests (black points) more visible.

```
par(mar = c(0, 0, 0, 0));
landscape    <- land_eg;
for(i in 1:dim(pop_last_time_step_sim1)[1]){
    xloc <- pop_last_time_step_sim1[i, 1] + 1;
    yloc <- pop_last_time_step_sim1[i, 2] + 1;
    landscape[xloc, yloc] <- 18;
}
land_cols <- c(land_cols, "#000000");
image(landscape, xaxt = "n", yaxt = "n", col = land_cols);
```

```
head(pop_last_time_step_sim1);
```

```
##    V3  V4     V102     V103      V104      V105
## 1  33  13 2.395557 3.485652 -1.413820  2.185108
## 2  94 109 4.325184 1.408115  1.212488 -0.505254
## 3 108  58 4.583821 3.336972  2.907757 -1.297925
## 4 108  61 3.385105 2.619580  1.399597 -0.577378
## 5   2  47 3.713334 3.623724 -0.853502  2.222031
## 6  87  63 4.034968 3.601765  1.555337 -0.060638
```

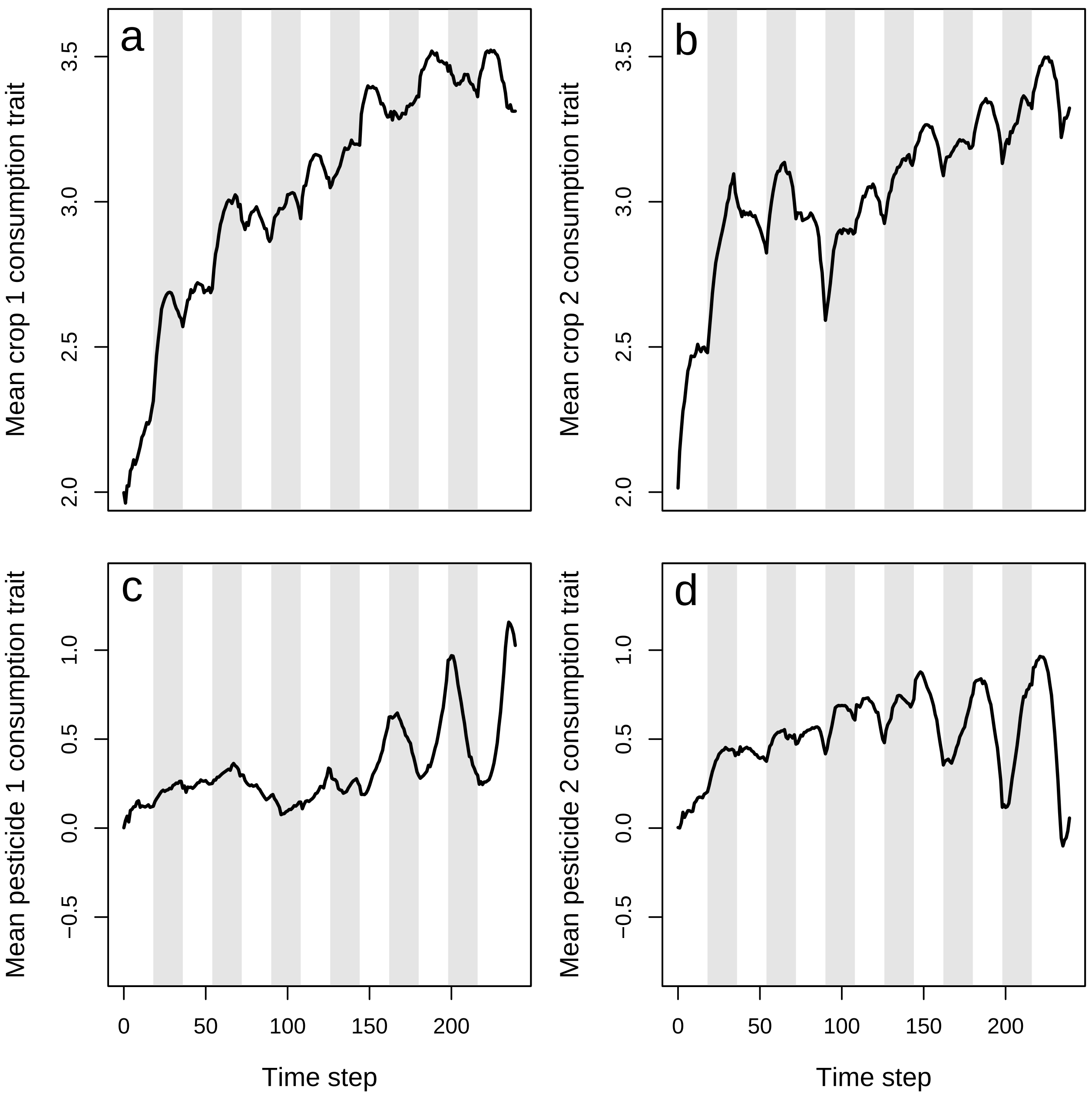


Figure 6: Mean values of pest traits T1 (crop 1 consumption rate), T2 (crop 2 consumption rate), T3 (pesticide 1 consumption rate), and T4 (pesticide 2 consumption rate) in a population simulated using the resevol R package in which pesticides are not rotated over time. Grey shaded regions show individual crop seasons; pesticides are applied at the start and midpoint of each seasons, beginning at the time step 100.

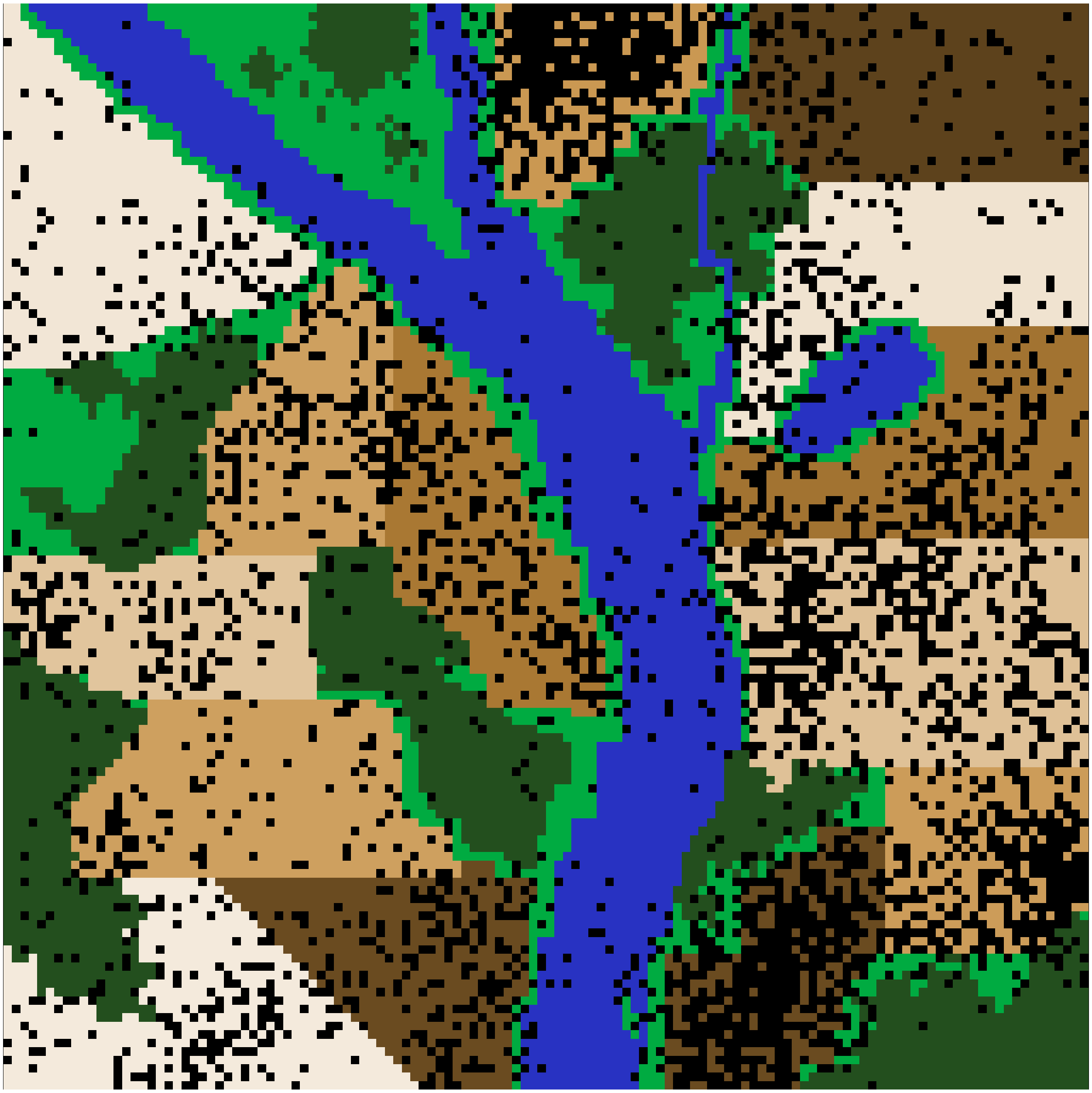

Figure 7: A complex landscape for simulating the ecology of pests and the evolution of pesticide resistance on farmland in the resevol R package. Terrain includes farms (brown colours), grassland (light green), forest (dark green), and water (blue). Black points show the locations of individual pests after 240 time steps for a simulation in which pesticides are not rotated over time.

```
print(land_cols);
```

```
##  [1] "#f4eadc" "#6a4b20" "#cea05f" "#e1c59d" "#a97833" "#cea05f" "#f2e6d6"
##  [8] "#6a4b20" "#cc9c59" "#dfc197" "#a27331" "#f0e3d0" "#5d421c" "#ca9852"
## [15] "#00ab41" "#234F1E" "#2832C2" "#000000"
```

The spatial pattern of pests over the landscape can be useful for visualising pest population dynamics. In the case of the above, pest density is higher on some farms (e.g., 1, 8, 9, 10, and 15) than others (e.g., 3, 12, and 13), and this pest density could be interpreted with respect to individual farm crop and pesticide rotation regimes. Hence, for the four evolving traits, a covariance matrix of pest traits can be calculated at the end of the simulation.

```
end_trait_covs <- cov(pop_last_time_step_sim1[,3:6]);
print(end_trait_covs);
```

```
##            V102       V103       V104       V105
## V102  1.1223660 -0.1485266  0.2470207  0.1506317
## V103 -0.1485266  1.3351424 -0.0714354  0.5612672
## V104  0.2470207 -0.0714354  2.9213039 -2.8459986
## V105  0.1506317  0.5612672 -2.8459986  3.1352726
```

The above covariance matrix shows that after 240 time steps, the covariance between consumption of pesticides 1 and 2 has decreased from -0.3992066 to -2.8459986, suggesting that the trade-off for resisting pesticides had become stronger. In contrast, the covariance between consumption of crops 1 and 2 has increased from -0.4510309 to -0.1485266, suggesting that the trade-off in ability to consume crops of each type has weakened.

#### 4.2 Simulation 2

We now repeat the simulations such that pesticide application is rotated every 9 time steps. To make this rotation, we only need to redefine `rotate_pesticide`.

```
rotate_pesticide           <- matrix(data = 0, nrow = 3, ncol = 3);
rotate_pesticide[1, 2]     <- 1;
rotate_pesticide[2, 1]     <- 1;
rotate_pesticide[3, 3]     <- 1;
```

Now, every 9 time steps, crops using pesticide 1 will switch to pesticide 2, and crops using pesticide 2 will switch to pesticide 1. We can re-run the same simulation with the same genome (`mg`), landscape (`land_eg`), and simulation seed (2022) used in simulation 1. Hence, the only thing changing between the two simulations is the pesticide rotation.

```
set.seed(2022);
sim2       <- run_farm_sim(mine_output             = mg,
                          terrain                  = land_eg,
                          crop_init                = initial_crop,
                          crop_rotation_type       = rotate_crop,
                          pesticide_init           = initial_pesticide,
                          pesticide_rotation_type  = rotate_pesticide,
                          food_consume             = c("T1", "T2", 0),
                          pesticide_consume        = c("T3", "T4", 0),
```

```
                                crop_number              = 3,
                                pesticide_number         = 3,
                                trait_means              = c(2, 2, 0.0, 0.0),
                                max_age                  = 6,
                                min_age_feed             = 0,
                                max_age_feed             = 2,
                                min_age_move             = 3,
                                max_age_move             = 6,
                                min_age_metabolism       = 3,
                                max_age_metabolism       = 6,
                                metabolism               = 0.5,
                                food_needed_surv         = 1,
                                reproduction_type        = "food_based",
                                food_needed_repr         = 2,
                                N                        = 1000,
                                repro                    = "biparental",
                                mating_distance          = 4,
                                rand_age                 = TRUE,
                                pesticide_tolerated_surv = 2,
                                movement_bouts           = 4,
                                move_distance            = 2,
                                crop_per_cell            = 10,
                                crop_sd                  = 0,
                                pesticide_per_cell       = 1,
                                pesticide_sd             = 0,
                                crop_rotation_time       = 18,
                                pesticide_rotation_time  = 9,
                                time_steps               = 240,
                                pesticide_start          = 81,
                                immigration_rate         = 100,
                                land_edge                = "reflect",
                                mutation_pr              = 0.01,
                                crossover_pr             = 0.01,
                                print_gens              = TRUE,
                                print_last              = TRUE);
```

We can look at the same outputs for simulation 2 that we did for simulation 1. We first read in the output for simulation 2.

```
population_data_file_sim2 <- system.file("population_data_sim2.csv",
                                         package = "resevol");
population_data_sim2      <- read.csv(file = population_data_file_sim2);
```

To avoid needless repetition, the code for plots will not be included for simulation 2. Figure 8 shows population abundance over time for simulation 2.

After pesticides are applied at time step 81, pest population size decreases and does not recover. Figure 9 shows crop and pesticide consumption over time.

Figure 10 shows how pest traits evolve over time when pesticides are rotated.

We can observe the spatial distribution of pests on the landscape in the final time step (Figure 11).

Finally, we can examine the trait covariances in the final time step below.

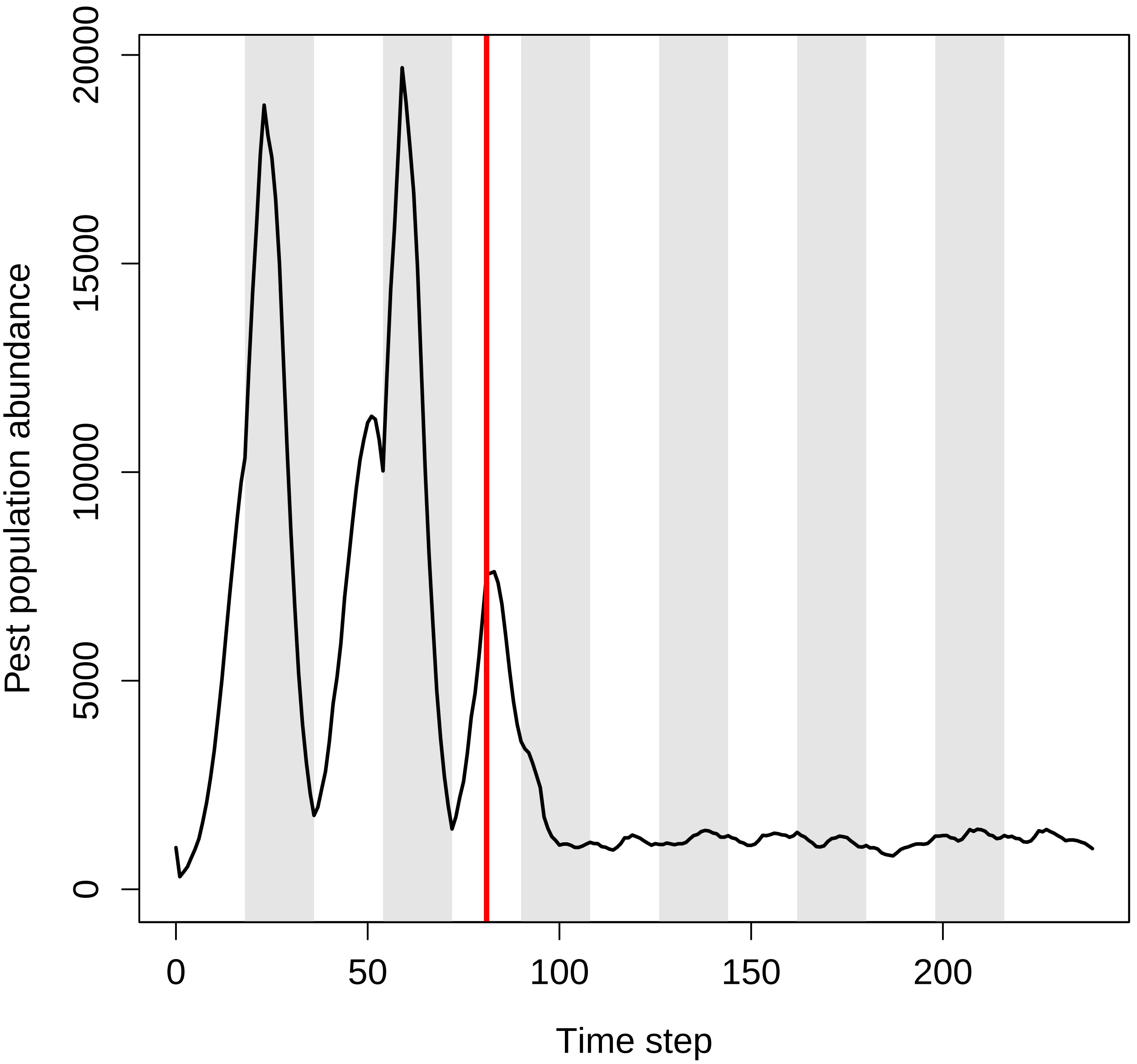


Figure 8: Pest abundance over time in a population simulated using the resevol R package in which pesticides rotates every 9 time steps. The grey shaded regions show individual crop seasons; pesticides are applied at the start and midpoint of each seasons, beginning at the time step indicated by the red vertical line.

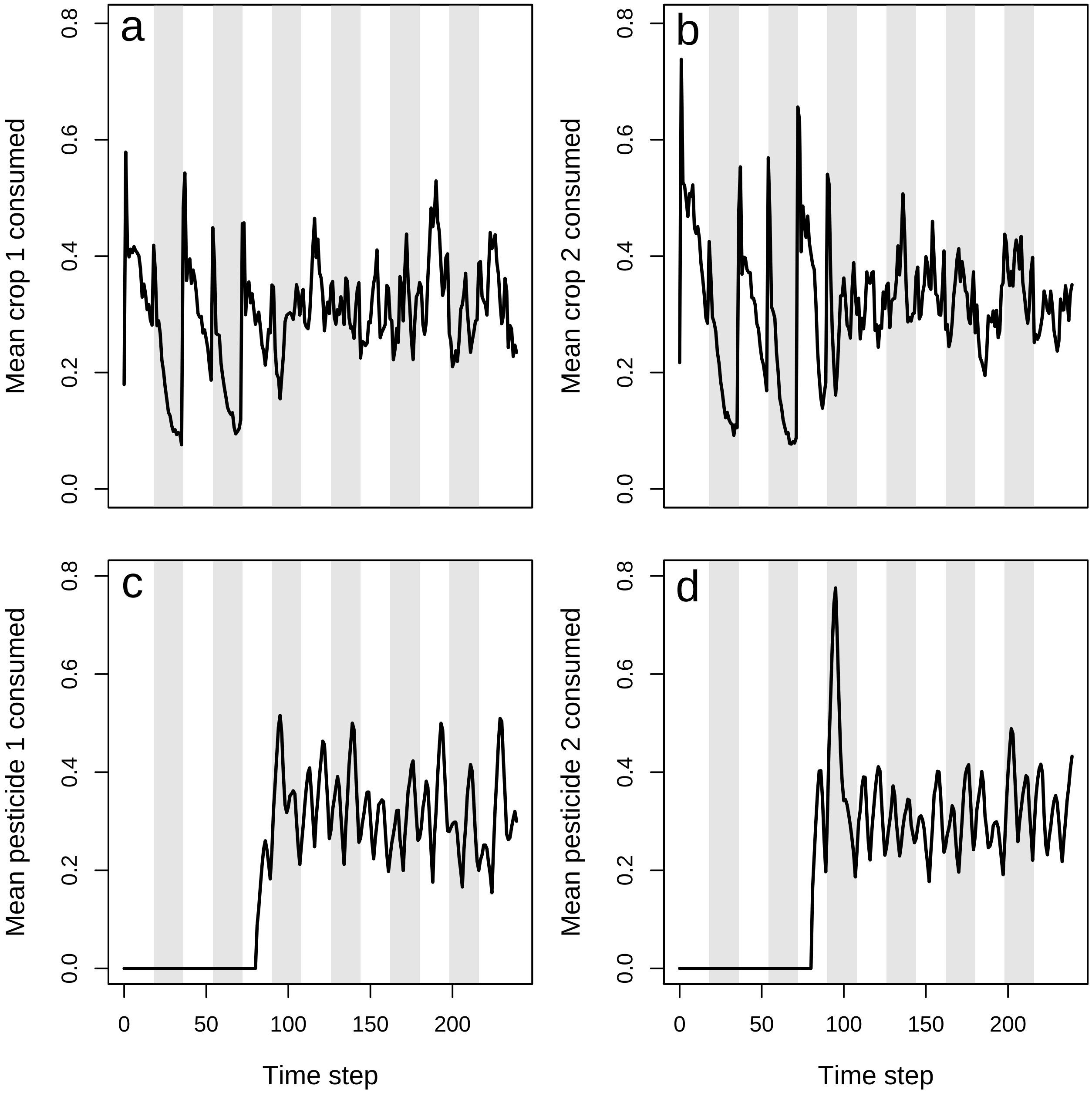


Figure 9: Pest consumption over time in a population simulated using the resevol R package in which pesticides rotate every 9 time steps. Panels show (a) mean amount of crop 1 and (b) crop 2 consumed per pest, and (c) mean amount of pesticide 1 and (d) pesticide 2 consumed per pest. Grey shaded regions show individual crop seasons; pesticides are applied at the start and midpoint of each seasons, beginning at the time step 100.

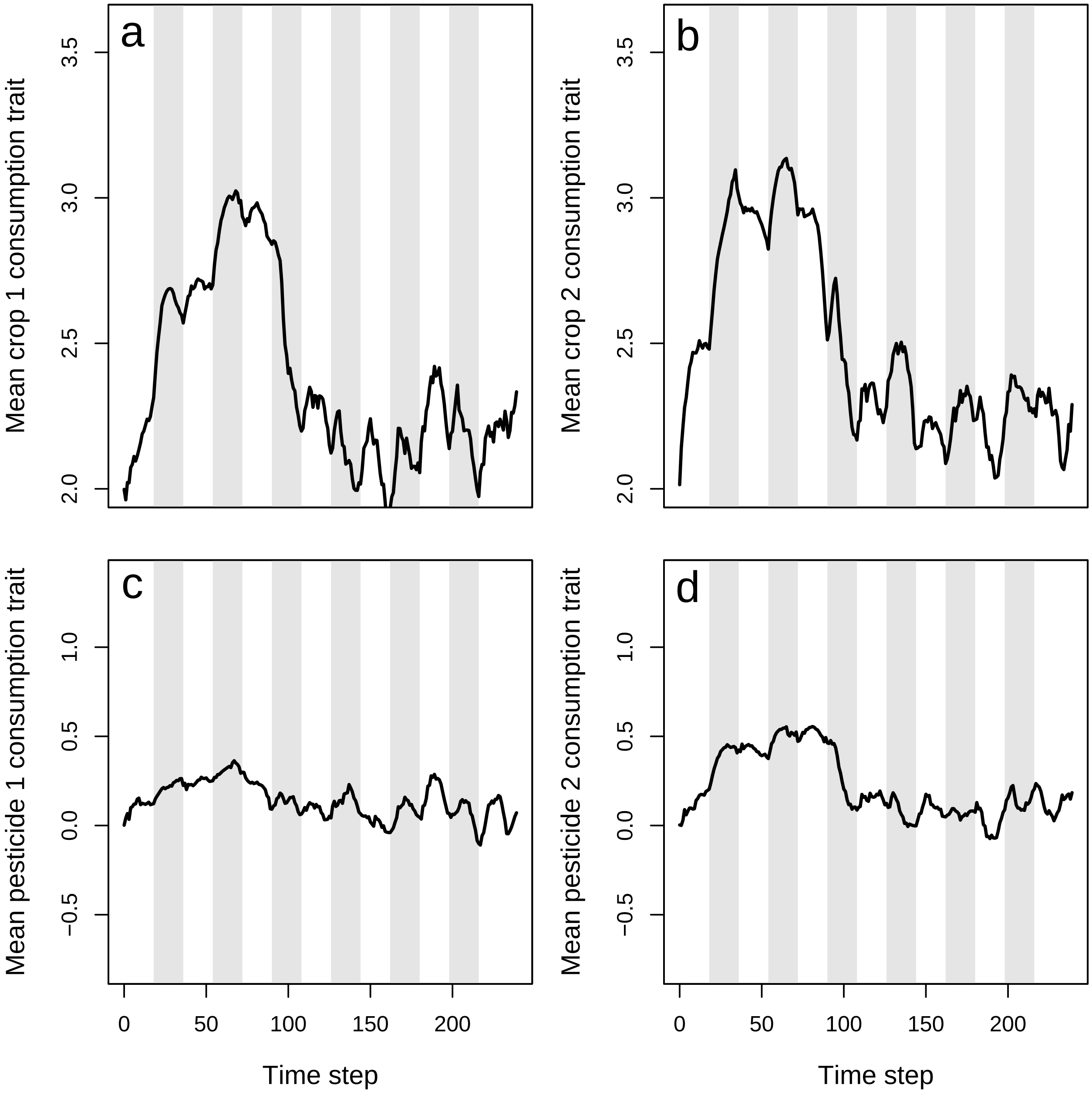


Figure 10: Mean values of pest traits T1 (crop 1 consumption rate), T2 (crop 2 consumption rate), T3 (pesticide 1 consumption rate), and T4 (pesticide 2 consumption rate) in a population simulated using the resevol R package in which pesticides rotate every 9 time steps. Grey shaded regions show individual crop seasons; pesticides are applied at the start and midpoint of each seasons, beginning at the time step 100.

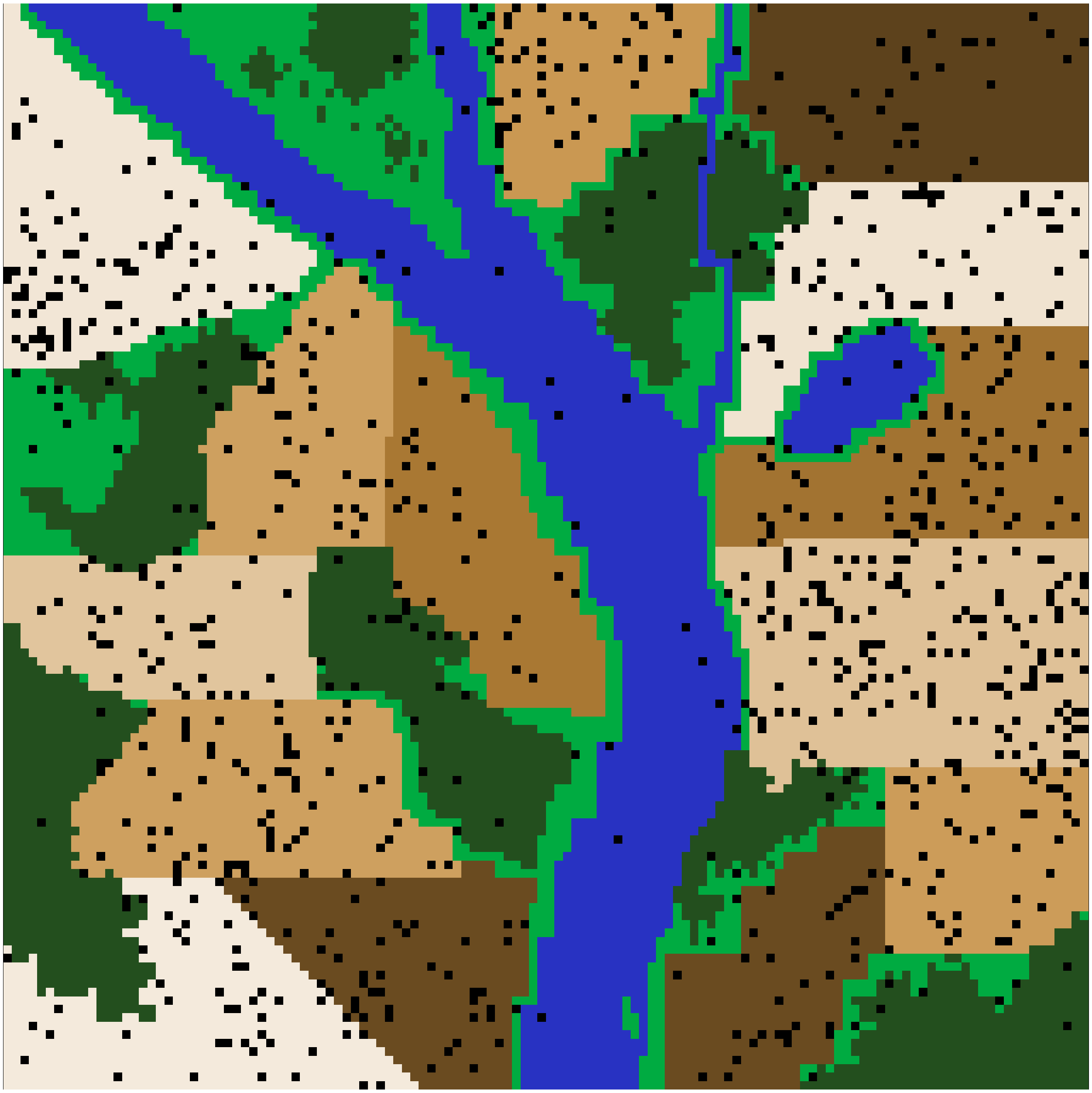

Figure 11: A complex landscape for simulating the ecology of pests and the evolution of pesticide resistance on farmland in the resevol R package. Terrain includes farms (brown colours), grassland (light green), forest (dark green), and water (blue). Black points show the locations of individual pests after 240 time steps for a simulation in which pesticides are not rotated over time.

```
##            V102        V103       V104        V105
## V102  1.5769217 -0.53211838  0.0665547  0.35835679
## V103 -0.5321184  1.29401550  0.2987174  0.01794992
## V104  0.0665547  0.29871744  0.9629416 -0.81135108
## V105  0.3583568  0.01794992 -0.8113511  0.96416686
```

We can use the outputs above to contrast the population dynamics of pests under a regime that lacks (simulation 1) or does not lack (simulation 2) the rotation of pesticide types.

### 5 Conclusion

From the output above, we can conclude that when farms rotate the pesticide that they apply between pesticide 1 and pesticide 2, it has a substantial effect on pest density and evolution. In the absence of pesticide rotation, the pest population quickly recovers after pesticides are first applied (Figure 4). Consequently, pesticides have very little long-term effect on the amount of crops consumed (Figure 5a,b), and while there is an initial spike in the consumption of both pesticides (Figure 5c,d), this consumption quickly declines. Despite the positive covariance between crop consumption and pesticide consumption, and the negative covariances between consumption of crop and pesticide types, these trade-offs appear to have only a modest effect on reducing actual crop consumption and maintaining or increasing pesticide consumption (Figure 6). Consequently, some farms experience high densities of pests that are resistant to the applied pesticides (Figure 7).

In contrast, when pesticides are rotated every 9 time steps, the pest population density drops after the onset of pesticide application and does not recover (Figure 8). The amount of each crop consumed per pests also drops (Figure 9a, b), while the amount of pesticide 1 and 2 consumed is maintained at a much higher rate than occurred in the absence of pesticide rotation (Figure 9c, d). Because selection against pesticide consumption was not consistent for a specific farm, and there is a trade-off between pest consumption of pesticide 1 versus 2, local adaptation to a single pesticide did not occur (Figure 10). Overall, farms therefore did not experience high densities of pests, and pest resistance to pesticide is well-managed (Figure 11).

Trait covariance differences between simulations 1 and 2 also highlight the effect that pesticide rotation had on pesticide resistance evolution. When no pesticide rotation occurred, the trade-off between consumption rate of pesticides 1 and 2 was reflected in a realised trait covariance of -2.8459986. But when pesticides were rotated, the realised covariance was instead -0.8113511. This means that in simulation 1, pests that are highly resistant to pesticide 1 were not highly resistant to pesticide 2, and vice versa. In other words, the lack of pesticide rotation resulted in local adaptation, with some pests specialising on resistance to one of the two pesticides. In simulation 2, pests maintained at least some resistance to both pesticides. Local adaptation was not possible because different pesticides were applied in sequence on any given farm.

This advanced example was not rigorous, but it illustrates how a rigorous simulation of pesticide resistance evolution could be designed with the resevol R package. To make robust predictions, multiple replicate simulations would need to be run for the same set of simulation conditions (in this case, for simulation 1 and 2) to account for stochastic effects in the model. For example, we might repeat the above initalisation of pest genomes to obtain 20 genomes with similar covariance structures, then run simulation 1 and simulation 2 20 times to get more robust predictions about pest population dynamics. We might also need to consider a range of parameter values if some values are unknown or uncertain (e.g., pest movement or trait means). To develop theory, we could also contrast simulations with more fundamental differences. For example, we might test how mating system affects pesticide resistance evolution by running the same set of simulations for `repro = "asexual"`, `repro = "sexual"`, and `repro = "biparental"`. The resevol R package is therefore a highly flexible and powerful tool for running complex models tailored to specific systems, and for developing theory on pest ecology, evolution, and management.
